# Proximal Tubule-on-Chip for Predicting Cation Transport: Dynamic Insights into Drug Transporter Expression and Function

**DOI:** 10.1101/2024.10.12.617976

**Authors:** Isy Petit, Quentin Faucher, Jean-Sébastien Bernard, Perrine Giunchi, Antoine Humeau, François-Ludovic Sauvage, Pierre Marquet, Nicolas Védrenne, Florent Di Meo

## Abstract

Deciphering the sources of variability in drug responses requires to understand the processes modulating drug pharmacokinetics. However, pharmacological research suffers from poor reproducibility across clinical, animal, and experimental models. Predictivity can be improved by using Organs-on-Chips, which are more physiological, human-oriented, micro-engineered devices that include microfluidics. OoC are particularly relevant at the fundamental and preclinical stages of drug development by providing more accurate assessment of key pharmacokinetic events. We have developed a proximal tubule-on-a-chip model combining commercial microfluidic and chip technologies. Using the RPTEC/TERT1 cell line, we set up a dual-flow system with antiparallel flows to mimic the dynamics of blood and urine. We assessed transporters mRNA expression using RT-qPCR, cellular polarization and protein expression via immunofluorescence and confocal microscopy, and monitored the transcellular transport of a list of prototypic xenobiotics by determining their efflux ratios with LC-MS/MS. Our results show that flow exposure significantly modulate mRNA expression of drug membrane transporters compared to static conditions. Dynamic conditions also enhance cell polarization, as evidenced by preferential basal and apical expressions of Na+/K+-ATPase, P-gp, OCT2, and MATE1, as well as the cellular secretory profile. We demonstrated unidirectional transcellular transport of a cationic substrate (metformin) with a higher efflux than influx ratio, inhibited with a specific OCT2 inhibitor, thus confirming the relevance of our proximal tubule-on-a-chip set up for cation transport investigations. Our proximal tubule-on-a-chip can also be used to explore the interactions between transporters, xenobiotics, and endogenous metabolites, possibly involved in the variability of individual drug responses. This study provides additional evidence that OoC can bridge the gaps between systemic and local pharmacokinetics, i.e., drug concentration close to its target, at the fundamental and preclinical stages.

**Highlights:**

- Cell exposure to flow shear stress modulate mRNA drug membrane transporters and cell polarization of proximal tubule
- Proximal tubule-on-chip relying on RPTEC/TERT1 cell line is a suitable platform for assessing transcellular cationic transport
- OCT2 and MATE are involved in potential drug-endogenous metabolite interactions
- Cell exposure to xenobiotics and endogenous metabolites modulate the drug transporters expression.

**Graphical Abstract:** 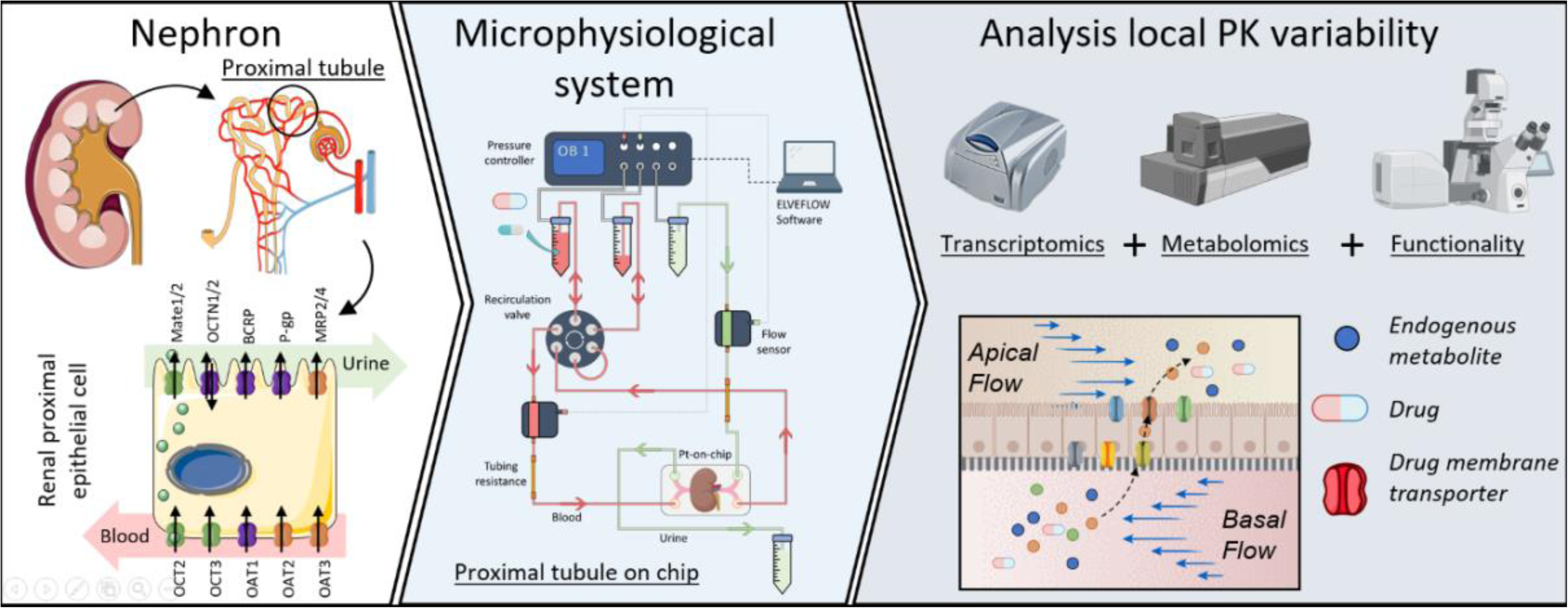

**Data Statement:** Original microscopy pictures, raw data for metabolomics and other data are available upon reasonable request.

## 1. Introduction

The inherent limitations of preclinical models drastically underestimate the high complexity of the human body, leading to discrepancies between observations obtained at different stages (clinical, *in vivo*, *in vitro* and *in silico*). Over the past decades, substantial efforts have been carried out to unravel pharmacological events governing xenobiotic concentrations (pharmacokinetics, PK) either locally or systematically^1,2^. In this context, drug transporters expressed at the intestinal, hepatic, renal, blood-brain and renal barriers, including the Solute Carrier (SLC) and ATP-binding cassette (ABC) families, are particularly relevant. This membrane proteins facilitate the transport of endogenous metabolites as well as xenobiotics from one compartment to another across epithelial barriers. Those expressed in the kidney proximal tubule (PT) play an important role in drug disposition. Since 2010, The International Transporter Consortium (ITC) has identified several drug membrane transporters of clinical importance that should or must be investigated not only in drug development but also retrospectively, after drug approval if not done before^3–7^. This list includes SLCs, namely organic anion transporters (*SLC22A6*/OAT1 and *SLC22A8*/OAT3), organic cation transporter 2 (*SLC22A2*/OCT2) and multidrug and toxin extrusion proteins (*SLC47A1*/MATE1 and *SLC47A2*/MATE2-K) as well as ABC transporters such as P-glycoprotein (*ABCB1*/P-gp). Importantly, these transporters are also cited by regulatory agencies (*e.g.*, the Food and Drug Administration and the European Medicines Agency) in their guidance documents for drug development, with a particular focus on drug-drug interactions (DDI). Noteworthy, retrospective investigations on multidrug resistance associated proteins (*ABCC2*/MRP2 and *ABCC4*/MRP4) are also recommended by ITC to elucidate their potential involvement in drug disposition from clinical observations^7^.

These PT transporters also participate in maintaining endogenous metabolite homeostasis^8,9^. Their physiological-pharmacological duality is now leveraged by the Remote Sensing and Signaling (RSS) Theory that relies on the presence of multi-specific and oligo-specific transporters in different organs that form a multi-scale, adaptive communication network that maintain or restore endogenous metabolite homeostasis^10^. One can easily hypothesize that drugs and their metabolite modulate the so-called RSS network via membrane transporters. Likewise, endogenous metabolite might influence local drug PK by affecting their intracellular and tissue concentrations. Such transporter-mediated drug-endogenous metabolite interactions (DMI) can represent a source of PK variability yet poorly explored, and dysregulate endogenous metabolism leading to off-target pharmacodynamics (PD) effects, locally as well as at distant sites.

Conventional models are not suited to study these interactions. On the one hand, differences between murine and human membrane transporters can lead to poor prediction of DDI and DMI^11,12^. On the other hand, conventional static 2D *in vitro* cell models do not sufficiently recapitulate the physiological or pathological mechanisms occurring *in situ*^13^. However, the last decade has seen the advent of new *in vitro* models called “microphysiological systems”, defined as devices or culture methods aimed at improving the physiological response of cells in experimental models. Organs-on-chips (OoCs) are defined as cell culture devices and methods that include a microfluidic-controlled microenvironment ^14,15^. These systems recreate physiological flows in the target organ, such as urine and blood flow in the case of renal PT. The flow is perceived by cells thanks to mechanosensory mechanisms^16^, including primary cilia, which in turn favor cell polarization, protein expression and metabolic functions^17–20^. One of the main challenges of OoC is achieving reproducibility, and another is standardization of protocols across laboratories^21,22^. The sources of variability are many, including the flow system setup, tubing length and diameter, chip geometry (often homemade), flowrate, exposure time, etc. Only a few teams provided detailed descriptions of their operating modes. Most published kidneys-on-chips used proximal tubule cells, as this part of the nephron is the main kidney site for drug reabsorption, secretion^23^ and a critical target for drug-induced nephrotoxicity^24–26^. The most commonly used kidney-on-chip configuration is an applied mono flow on tubular epithelial cells culture on 2D^26^. Additionally, only a few groups have studied the unidirectional transcellular transport of xenobiotics, and/or DDI and DMI. Noteworthy, Ma et *al.* have recently proposed a PT-on-chip based on induced human stem cells and a homemade device for functional investigations of renal transporters^27^. However, such models are challenging to scale up for drug screening due to their complexity (cellular heterogeneity and lack of maturation), moreover most do not use commercial devices but custom-made chips. The present study aimed to develop a PT-on-chip model for the purpose of investigating DDI and DMI as a potential source of local PK variability, as the first step towards the development of a platform encompassing several key pharmacological organs for systemic pharmacology purposes. It relies on the use of commercially available immortalized RPTEC/TERT1 cell lines, micro-perfusion system and chips. Given the well-known poor expression of OAT transporters in RPTEC/TERT1 cells^28,29^, we focused on the interplay between cationic probes and the OCT2/MATEs-driven transcellular transport, accounting for DDI and DMI. Using metformin and creatinine as pharmacological and endogenous probes respectively, we confirmed our PT-on-chip is functional and, has pharmacological relevance. Transporter-mediated DDI or DMI through the unidirectional OCT2/MATE transcellular transport were further explored considering transport inhibitors.

## 2. Materials and methods

### 2.1 Cell culture

RPTEC/TERT1 were obtained from American Type Culture Collection (ATCC) (ATCC, reference: CRL-4031), and cultured in Dulbecco’s modified Eagle’s medium with Ham’s F-12 nutrient mix (DMEM: F12) (ATCC, reference: 30-2006) supplemented with hTERT immortalized RPTEC Growth kit (ATCC, reference: ACS-4007), penicillin/streptomycin (Gibco, reference: 15140122) at 1% and Geneticin (Gibco, reference: 10131035) at 0.2%. No FBS was used. Cells were cultured routinely in T75 flasks (Sarstedt, TC Flask T75, Cell+, Vented Cap, reference: 83.3911.302) at 37°C in a 5% CO2 humidified atmosphere. Cells were sub cultured by trypsinization with 0.25% trypsin (Gibco, reference: 25200056), and the media was changed every two days. The cells used for further experiments were at passages 4 to 15.

### 2.2 Microfluidics setup

The Elveflow OB1 MK4 microfluidic system from Elveflow (Paris, France) (Figure S1) include a pressure- and flow-controller that apply pressure on a reservoir connected to a microfluidic flow sensor. It adapts the pressure to maintain the user-defined flowrate. A recirculation pump was added to the system to limit media consumption and above all to prolong the contact time between cells and fluids. The tubing setups (lengths, diameter) are reported in Figure S1.

### 2.3 Proximal tubule-on-a-chip

RPTEC/TERT1 were seeded at 130 000 cells/cm² on µ-slide I luer ibitreat chips (ibidi, Cat. No: 80176) or on Be-doubleflow standard chips (Beonchip), depending on the experiment. All supports were thin coated with collagen 1 (Gibco, reference: A1048301) at 5 µg/cm² according to the manufacturer’s protocol. µ-Slide I luer ibitreat consists in one channel of 50 mm length, 5 mm width and 0.4 mm height. Be-doubleflow chip consists in 2 channels of 46 mm length, 1.5 mm width and 0.375 mm height. Channels are separated by a semi-porous membrane with a random distribution of pores across an average diameter of 1µM. Six hours after seeding, cell cultures on chips were placed on a cell culture rocker with a 5% angle. Cells were cultured for 10 days at 37°C in a 5% CO_2_ humidified atmosphere, and the culture medium was changed every day.

### 2.4 Quantification of mRNA transcripts

NucleoSpin RNA (Macherey-nagel, reference: 740955.250) or NucleoSpin RNA XS (Macherey-nagel, reference: 740902.50) kits was used to extract mRNA, according to the manufacturer’s protocols. Complementary DNA was obtained using the Master Mix SuperScript™ IV VILO™ (reference:11756050), according to the manufacturer’s protocol. Gene-specific primer-probe (table S1) sets and Master Mix TaqMan™ Fast Advanced, no UNG (reference: A44360) was purchased from ThermoFisher scientific. Quantitative PCR (qPCR) reactions were carried out using the Rotor-Gene Q 2plex from QIAGEN. Gene expression levels were normalized to GAPDH expression and expressed as fold-difference compared to the control.

### 2.5 Immunofluorescent cell staining for confocal imaging

For immunofluorescent staining, cells were fixed with 1% PFA for 10 min at room temperature. After 3 steps of washing with phosphate-buffered saline (PBS, Gibco, reference: 14190-094), the non-specific sites were saturated with 3% PBS-Bovine serum albumin (BSA, Sigma, reference: A9418-10G) for 1 hour. After washing steps with PBS, the chips were spiked with a primary antibody diluted with PBS-BSA 3% for 12 h at 4°C. After another 3 PBS washing steps, chips were treated with a solution of secondary antibodies (Table S2) for 1 h at room temperature. After PBS washing, chips were spiked with DAPI (Invitrogen, reference: D1821) at [300 nM] for 5 min. After PBS washing, the chips were spiked with the CitiFluor^TM^ AF1 mountant solution (Electron microscopy sciences, cat. N° 17970.25) before confocal imaging with ZEISS LSM880.

### 2.6 Polarized transport study

#### 2.6.1 Transporter activity assay

RPTEC/TERT1 were seeded on the apical channel of BE double flow chip, and cultured as described previously, after ten days they were put under antiparallel flows at 10 µL/min (or 0.03 dyn/cm²) for the apical channel to model urine flow and 20 µL/min (0.07 dyn/cm²) for the basal chanal to model blood flow (Figure S1). After 24h under flow transcellular transport through the PT epithelial barrier was evaluated by measuring the apparent permeability for metformin [100 µM] (M), creatinine [10 µM] (C) or tenofovir [30µM] (T) at the apical (A) or basal (B) sides, for A-to-B or B-to-A permeability assessment 24h later. Permeability was also measured after transport inhibition with, cimetidine [100µM], ritonavir [15µM] and diclofenac [30µM]. Circulating fluids were collected at the beginning (T_0_) and after 24 hours of substrate exposure. Apparent permeabilities were measured under both the undetermined flow on a cell culture rocker or under flow shear stress (FSS) conditions .

#### 2.6.2 Compound quantitation by LC-MS/MS

The extraction of extracellular metabolites was performed as already described in Faucher et *al*^30^. The cell culture medium was collected and centrifuged at 3000 g for 1 min at room temperature to eliminate cell debris. Then 200 μL of acetonitrile and 20 μL of internal standard solution (2-isopropylmalic acid, 0.5 mM) were added to 100 μL of supernatant. Solutions were homogenized by vortex mixing for 30 s and centrifuged at 15000 g for 15 min at room temperature. Solutions were diluted 1/10 with ultrapure water and transferred into an injection vial for mass spectrometry analysis. Three microliters of the extracts were injected into the LCMS-8060 (Shimadzu, Kyoto, Japan) liquid chromatography - tandem mass spectrometry system. Metformin, tenofovir and creatinine concentrations were measured against a calibration range.

#### 2.6.3 Efflux Ratio calculation

The concentration obtained by mass spectrometry were used to calculate the efflux ratio, as follows:

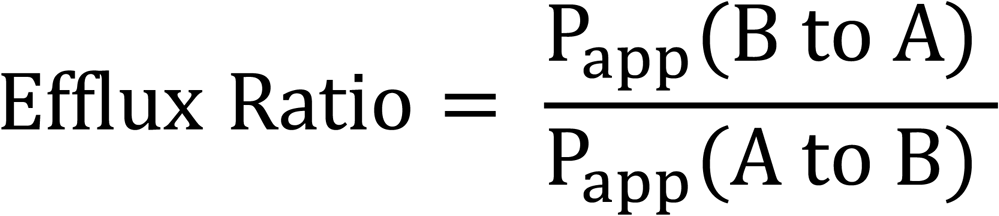

where P_app_ is the apparent permeability coefficient (cm/s) calculated for each compound for the apical-to-basolateral (A to B) and basolateral-to-apical (B to A) directed transport:

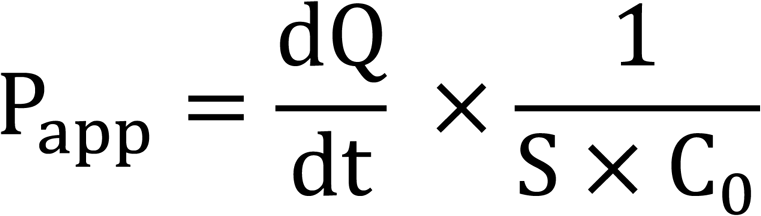

where 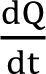 is the rate of drug appearance in the receiver compartment (mol/s), S is the surface area of the membrane (cm²), C_0_ is the initial concentration of the drug in the donor compartment (mol/cm³).

### 2.7 Metabolomics study

Extracellular metabolites were extracted from the cell culture medium as described in section 2.6.2. three milliliters of the extracts were injected into the same analytical system as above, using the LC-MS/MS “Method Package for Cell Culture Profiling Ver.2” (Shimadzu) where mass transitions of additional compounds had been added locally by infusing the pure substances in the mass spectrometer. For each transition analyzed, only well-defined chromatographic peaks were considered. The area under the curve of each metabolite was normalized to the area under the curve of the internal standard (2-isopropylmalic acid).

### 2.8 Statistical analysis

One-way analyses of variance (ANOVA) with Dunnett multiple comparison tests, or Kruskal-Wallis tests were performed using GraphPad (GraphPad Software Inc., San Diego, CA, USA). All data are presented as means +/- standard deviation; differences between groups were considered statistically significant for p < 0.05.

## 3. Results

### 3.1. Impact of flow exposure on cell transporter expression and metabolome

#### 3.1.1. mRNA expression of drug membrane transporters

Given the inherent challenges regarding reproducibility and harmonization across OoC setups, the RPTEC-TERT1 cell culture protocol was first optimized as described in Figure 1A. The best compromise between cell viability, monolayer architecture and transporter expressions was achieved by: (i) seeding at 130 000 cells/cm^2^; (ii) exposing post-confluence monocellular layers to undetermined flow for 9 days using a rocker; and (iii) finally applying predefined FSS for 4 days. In the present section, we employed commercial mono-channel chips to investigate the effects of flow.

**Figure 1.**
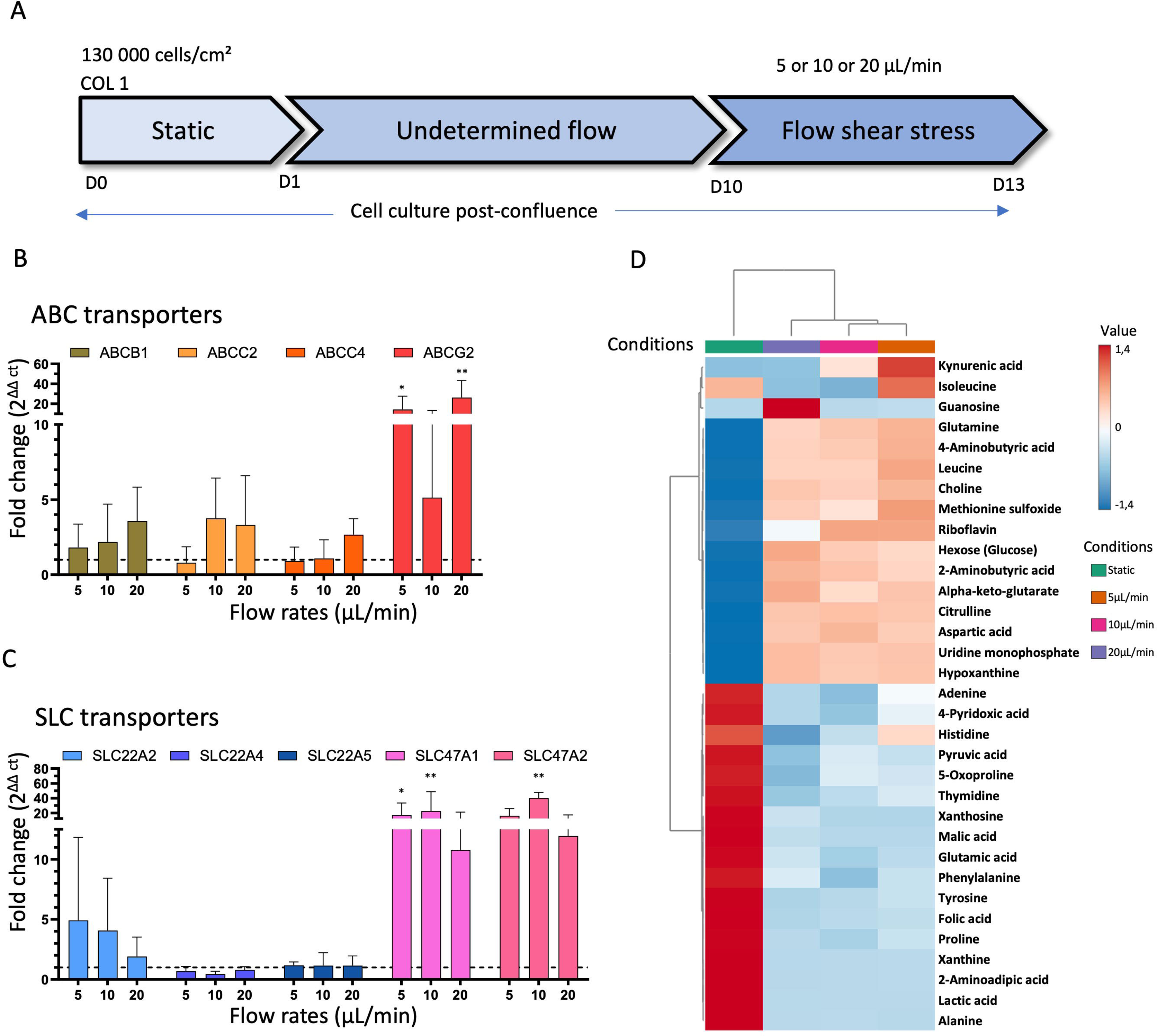
Flow shear stress influences drug transporter transcription and endogenous metabolite expression. (a) Protocol used for RPTEC/TERT1 cell culture in proximal tubule-on-chip in the present study. (b) Fold changes regarding the mRNA expressions of proximal tubule ABC and SLC transporters of pharmacological relevance. Bar plots show relative expression levels of RPTEC/TERT1 mRNA transporter expressions using mono-channel device where cells are exposed to different flow rates, 5 µL/min (0,0045 dyn/cm², N=6), 10 µL/min (0,009 dyn/cm², N=11) and 20 µL/min (0,02 dyn/cm², N=4). It is worth mentioning that for *SLC22A4*, *SLC22A5* and *SLC47A2*, only three independent replicas (N=3) were performed. Expression levels were normalized to the internal control GAPDH gene and are presented as fold changes relative to static conditions (N=5, dotted line). Error bars represent standard deviations. Statistical significances were conducting using one-way ANOVA analysis, asterisks (*) indicate statistically significant differences compared to static condition (*p-value < 0.05, **p-value < 0.01, ***p-value < 0.001). (c) Targeted search for 33 endogenous metabolites out of 103 initially considered (see main text) by LC-MS/MS on extracellular supernatant of RPTEC/TERT1 cells culture in static condition or exposed to different flow rates (5, 10 and 20μL/min). Data were standardized by auto scaling and hierarchically clustered with Ward method and represented by mean of 3 independent replicates. Metabolomic clustering was achieved using the MetaboAnalyst 6.0 Statistical Analysis online tool.

Different flow rates (5, 10 and 20 μL/min *i.e.*, 0.0045, 0.009 and 0.02 dyn/cm^2^, respectively) were first considered to assess the mRNA expression of drug membrane transporters of pharmacological relevance. Particular attention was paid to *ABCB1*, *ABCC2*, *ABCC4*, *ABCG2* (see Figure 1B) and *SLC22A2*, *SLC22A4*, *SLC22A5*, *SLC47A1* and *SLC47A2* (see Figure 1C). Controlled flow exposure was associated with increased mRNA expression of ABC transporters. Interestingly, the higher the flow rate, the higher the mRNA expression of *ABCB1*, *ABCC2* and *ABCC4*. By contrast, *ABCG2* gene expression was only significantly increased at 5 (p = 0.027) and 20 (p= 0.003) μL/min, but not at the intermediate 10 μL/min flow rate. Shear stress also increased the transcriptomic expression of SLC transporters, except for *SLC22A4* and *SLC22A5*, for which mRNA expression was either unchanged (*SLC22A4*) or decreased (*SLC22A5*, see Figure 1C). *SLC47A1* and *SLC47A2* were significantly increased at 5µL/min for SLC47A1 (p= 0.037) and at 10µL/min for SLC47A1 (p= .005) and SLC47A2 (p= 0.002). However, there are more sensitive to shear stress than *SLC22A2*. SLC transporters mRNA expression was not correlated with the flow rate. Additionally, mRNAs of *SLC22A6* nor *SLC22A8* were not detected under undetermined flow or FSS conditions, as expected with RPTEC/TERT1 cell lines^28,29^.

#### 3.1.2. Cell secretory profiles under flow conditions

Likewise, the cell excreted secretome was compared between undetermined flow and FSS conditions on commercial mono-channel chips, by monitoring metabolite concentrations in the medium. Serum-free conditions were used to ensure that the metabolites identified were not artefactual. More than 103 metabolites were searched (Table S3), of which only 33 were detected (Table S4). Interestingly, two different behaviors were observed: 13 metabolites exhibited increased concentrations and 17 decreased concentrations under flow conditions, as compared to the undetermined flow conditions (Figure 1D). Even though these modifications are mild, they highlight that shear stress modulates intracellular metabolisms and cellular absorption/secretion behaviors. Metabolic profiles did not exhibit a flow rate/concentration relationship likely due to experimental conditions, except for kynurenate and isoleucine. Thereby, no specific pathway can be identified, requiring further investigations which are out of the scope of the present study.

#### 3.1.3. Cell polarization and architecture

An epithelial barrier suited to model PT elimination function requires proper cell polarization, which is not often checked in conventional static 2D culture models^20^. We investigated cell polarization under dynamic conditions by monitoring protein expression and localization using immunofluorescence. We first focused on P-gp and Na+/K+-ATPase protein expression, indicating proper basal to apical polarization (Figure 2A). They were mostly expressed at the basal and apical membrane, respectively, as evidenced by z-stack analyses using confocal microscopy. Likewise, but to a lesser extent, OCT2 and MATE1 protein were also expressed at proper location, at the basal and apical membrane respectively (Figure 2A). Interestingly, immunofluorescence also indicated intracellular expressions of membrane transporters, but not Na+/K+-ATPase. Importantly, OAT1 was not observed by immunofluorescence, as expected from (i) the literature and (ii) their mRNA expression levels (see above). OAT3 was observed using immunostaining under the undetermined flow and FSS conditions (Figure S2).

**Figure 2.**
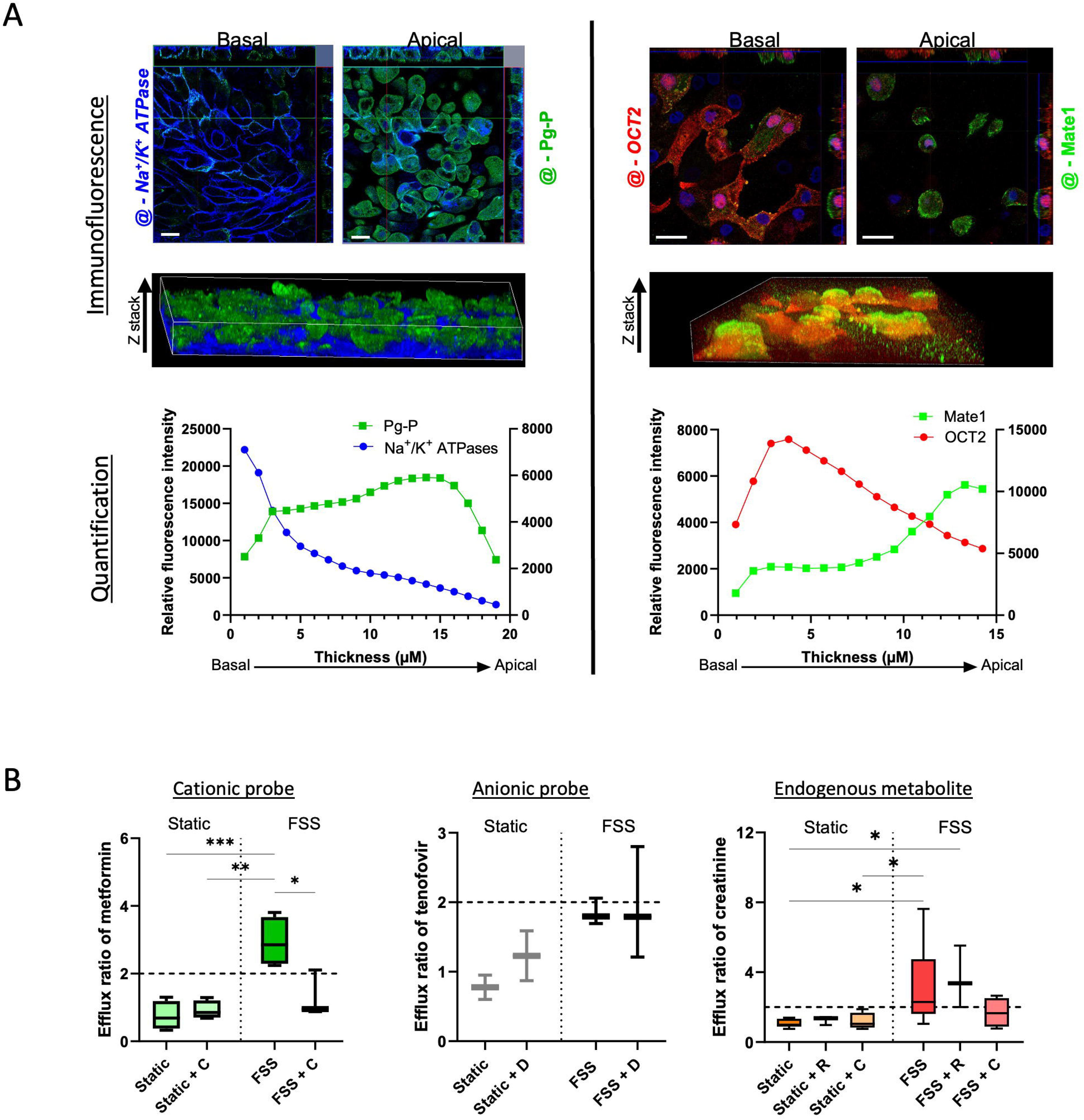
Expression and localization of specific transporter under flow conditions. (a) Apical and basal immunostaining of Na^+^/K^+^ ATPase (left panel, blue), Pg-P (left panel, green), OCT2 (right panel, red) and, MATE1 (right panel, green) conducted on RPTEC/TERT1 cell line in mono channel device under FSS (20µL/min, i.e., 0,02 dyn/cm²) as well as 3D reconstruction and quantification of relative fluorescent intensity along cell layer thickness. (b) Calculated efflux ratios of metformin (left), creatinine (center) and tenofovir (right) under undetermined flow and FSS. Experiments were either conducted with standalone substrate (metformin, 100µM, N=4; creatinine, 10μM, N=8; tenofovir, 30 μM, N = 3) or in presence of inhibitor (metformin/cimetidine, 100μM/100μM, N=3; creatinine/cimetidine, 10μM/100μM, N=4; creatinine/ritonavir, 10mM/15mM, N=3; and tenofovir/diclofenac, 30μM/30μM, N=4). Statistical significance was determined using one-way ANOVA, asterisks (*) indicate statistically significant differences compared with undetermined flow (*p-value < 0.05, **p- value < 0.01, ***p-value < 0.001).

### 3.2. Modeling xenobiotic elimination using PT-on-chip

#### 3.2.1. Transporter-mediated transcellular transport of pharmacological probes

Transporter-mediated transcellular transport was then investigated to ensure that the PT-on-chip setup developed reliably model one of the main pharmacological functions of PT epithelial cells, i.e., elimination of xenobiotics and metabolites from blood into urine. We check the relevance of dual channel chips as compared to mono channel devices by measure mRNA expression of membrane transporters (Figure S3). The epithelial barrier integrity was assessed by monitoring Dextran-FITC transport from the apical to the basal channel (Figure S4) were no impact of metformin treatment was detected.

We measured apparent permeabilities of metformin and creatinine, known substrate for the basal-apical OCT2-MATE transporter pair. We also perform assay for tenofovir as a negative control, since it is transported by OAT1/3 and MRP2/MRP4 at the basal and apical membranes, respectively (Figure S5). Interestingly, apparent permeabilities under FSS conditions are one order of magnitude lower than under undetermined flow condition. This might be explained by a lower substrate residence time due to higher flow velocity.

Efflux ratios were then calculated to measure the unidirectional (or vectorial) transcellular transport. For the OCT2/MATEs transporter pair, the vectorial transports under FSS were significantly higher for metformin (p< 0.001) and creatinine (p= 0.01) (efflux ratio: 2.94 ± 0.727 and 3.24 ± 2.21, respectively) (Figure 2B), than under undetermined flow, where no vectorial transcellular transport was observed (efflux ratio: 0.75 ± 0.42 and 1.06 ± 0.23, respectively). As expected, our PT-on-chip setup does not properly express vectorial transport involving OATs/MRPs pairs (tenofovir efflux ratio: 1.85 ± 0.19 and 0.78 ± 0.25 under the FSS and undetermined flow conditions, respectively), due to insufficient expression of influx OATs at the RPTEC/TERT basal membrane.

#### 3.2.2. Modeling transporter-mediated drug-drug and drug-metabolite interactions

We used cimetidine, a known OCT2/MATEs inhibitor, to investigate the pharmacological relevance of the present PT-on-chip model further. Under undetermined flow conditions, cimetidine did not influence metformin (exogenous substrate) and creatinine (endogenous substrate transport (efflux ratios: 0.92 ± 0.27 and 1.18 ± 0.43, respectively) (Figure 2B). In contrast, under FSS it decreased metformin (p= 0.016) and creatinine efflux (ratio: 1.31 ± 0.69 and 1.68 ± 0.85, respectively), supporting the inhibitory effect of cimetidine on either OCT2 or MATE1/MATE2K transporters. Creatinine efflux ratio was not significantly inhibited by ritonavir, a known inhibitor of OCT2 but not MATE (efflux ratio: 3.63 ± 1.77), suggesting a higher affinity of creatinine to OCT2 than ritonavir.

For the sake of comparison, an OAT inhibitor (diclofenac) was also associated with tenofovir, showing no effect on its efflux ratio (1.93 ± 0.81), but the latter was already below the threshold of 2 without inhibitor.

### 3.3. Modulating transporter expressions and secretome by transporter substrates

Under FSS, PT transporter expression and intracellular metabolism were investigated after xenobiotic or creatinine exposure. In the presence of metformin, transcriptomic analyses revealed an upward trend for mRNA expression of ABC and SLC transporters (significant for *ABCC2* (p=0.007), *ABCC4* (p=0.028), *SLC47A1* (p=0.012), *SLC47A2* (p=0.007) and *SLC22A5* (p=0.007)), except for *ABCG2* (Figure 3A). Interestingly, when cimetidine was added, mRNA expression increases were all abolished, except for *SLC47A1* whose mRNA expression was slightly, although not significantly, increased as compared to the control.

**Figure 3.**
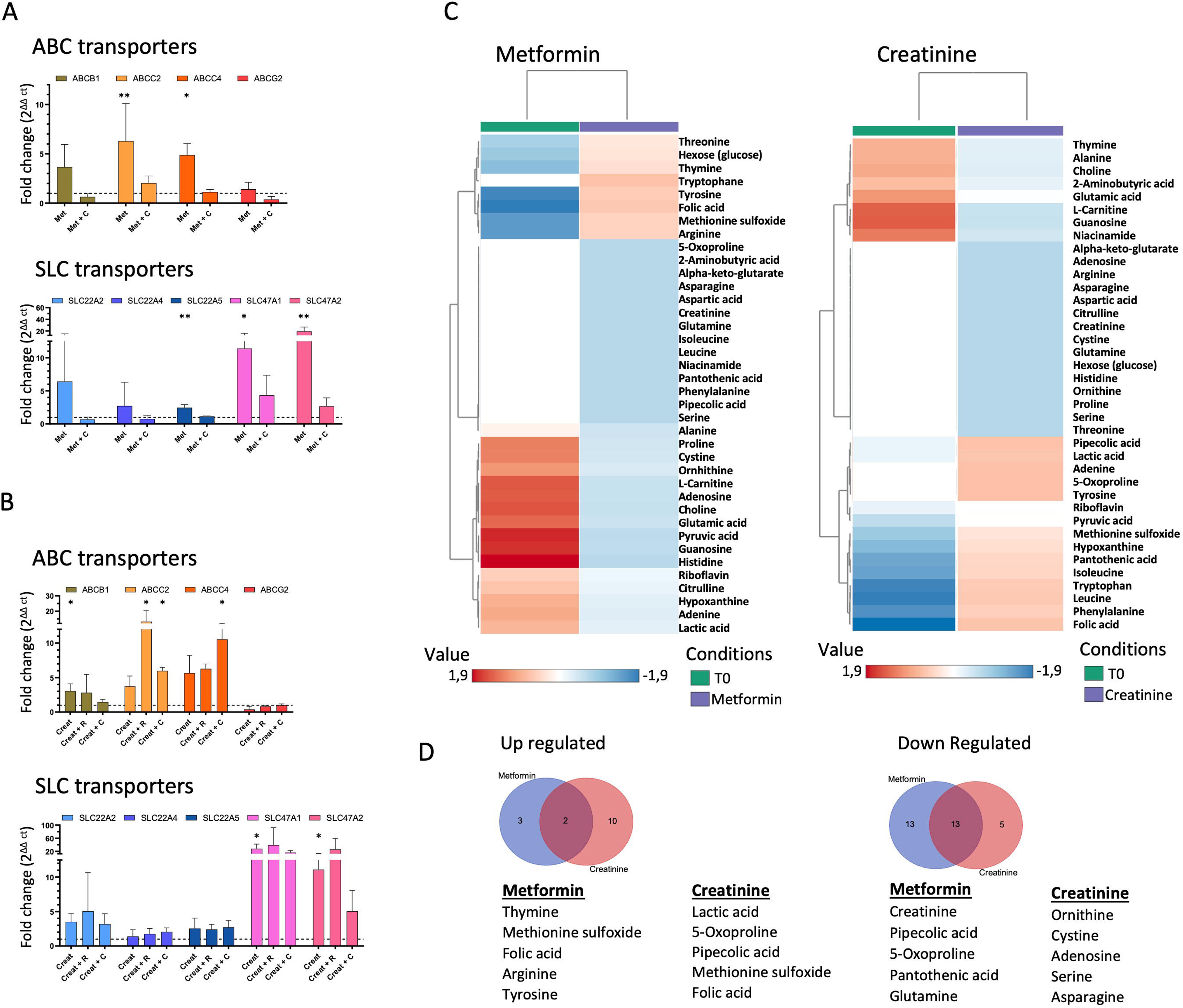
Metformin or creatinine treatments alter drug transporter transcriptional expression and endogenous metabolite expression. Fold changes regarding the relative expression levels of mRNA transporter expressions of RPTEC/TERT1 cell line in dual-channel device under treatment of (a) metformin (Met, 100µM, N=3), in presence or not of cimetidine (C, 100µM, N=3) and (b) creatinine (Creat, 10µM, N=3) in presence or not of ritonavir (R, 15µM, N=3) or cimetidine (C, 100µM, N=3). Error bars represent standard deviations. Statistical significance was determined using Kruskal Wallis test, asterisks (*) indicate statistically significant differences with respect to untreated PT-on-chip (N=5, *p-value < 0.05, **p-value < 0.01, ***p-value < 0.001). (b) Heat map of intracellular metabolites altered under metformin (left panel) creatinine (right panel) treatment. (c) Venn diagram of shared up regulated and down regulated metabolite under treatment including the top 5 most up and down regulated for each treatment separately.

In contrast, when PT-on-chip was exposed to creatinine, mRNA expression of most membrane transporters remained unchanged, whereas *ABCB1* (p=0.015), *SLC47A1* (p=0.032), *SLC47A2* (p=0.035) were significantly increased (Figure 3B). Adding ritonavir or cimetidine had no strong impact, except for *ABCC2 (*with cimetidine p=0.044; with ritonavir p=0.023) and *ABCC4 (*with cimetidine p=0.021) where an increase in expression was observed.

Intracellular metabolism was next evaluated under transporter substrate exposure and compared with the initial conditions T_0_ (Figure 3C). We identified 38 differentially expressed metabolites (Table S5). Five of them were up-regulated and 26 down-regulated by metformin. Creatinine exposure was associated with 12 and 18 up- and down-regulated metabolites, respectively (Figure 3D). Folic acid and methionine sulfoxide were modulated by both creatinine and metformin, while 13 other metabolites were down-regulated by both.

## 4. Discussion

We report here the development of a functional PT-on-chip set-up that model the transporter-mediated transcellular transport of cationic substrates, whether endogenous or exogenous. The influence of different FSSs on drug membrane transporter expression and intracellular metabolism was investigated to determine the optimal parameters of this model. FSS-dependent profiles regarding mRNA expression and transport functions suggest different responses among pharmacologically relevant drug membrane transporters. However, such a statement must be smoothed because of the inherent nature of the immortalized cell lines used which are known not to express all membrane transporters. Our results are consistent with former investigations^20,31,32^ supporting that dynamic microfluidic environments improve transporter expression and tubular reabsorption and secretion. For instance, flow-induced FSS increased the expression of the SGLT1 transporter and transepithelial glucose transport by primary human proximal tubular epithelial cells. Also, the efflux capacity of ABCB1/P-gp in the presence of FSS is stronger than in static conditions.^19^ Interestingly, several studies employed PT-on-chips to investigate pharmacologically relevant events. Vriend *et al.* developed the so-called Nephroscreen platform relying on the culture of immortalized human tubular cells. This high-throughput PT-on-chip system is applicable to xenobiotic screening to predict drug-induced nephrotoxicity.^33^

However, only a few *in vitro* models have been developed to mimic xenobiotic elimination through the tubular epithelial barrier. The present results show that our PT-on-chip setup of immortalized RPTEC/TERT1 cells and commercial chips recapitulates: (i) the unidirectional (or vectorial) transcellular transport of xenobiotics and endogenous metabolites through OCT2, MATE1 and MATE2-K transporters; and (ii) transporter mediated drug-drug or drug-metabolite interactions. Unfortunately, it cannot be used in its present form to model the unidirectional transcellular transport involving OATs since they are not expressed by the RPTEC cell line, as described in the literature. Only one very recently reported PT-on-chip setup, using demanding human induced pluripotent stem cells, recapitulate the elimination of anionic and cationic substrates^27^.

The organ-on-chip technology still suffers from a lack of reproducibility and transferability, due to differences in lab practices, as well as to inherent experimental variability (flow, tubing setup).^34,35^. Since 2015, the number of papers published is correlated with that of commercial solutions^26^. It is important to distinguish turnkey systems combining microfluidic platforms and integrated chips^36,37^ from adaptative setups including independent microfluidic systems and commercially available chips, tailored to the tissues and objectives. The former limit the integration of additional biological compartments in a physiologically relevant configuration, or accessories such as oxygen sensors or electrodes for barrier integrity measurements. We here propose a robust combination of commercial systems for microfluidic control and microchips, suited to our rather complex double-flow PT model.

With this experimental setup, our objective is to investigate further the underrated transported- mediated interplay between xenobiotics and endogenous metabolites. This new concept, named “remote sensing signaling (RSS) theory”, was originally defined as hormone-free inter-organ communication ensuring the homeostasis of small endogenous metabolites^10,38–40^. It has been recently proposed that key pharmacological partners, such as membrane transporters and drug metabolism enzymes, might play a role in the so-called RSS network. Therefore, our hypothesis is that drug-metabolite interactions, though RSS, may participate in the intra- and inter-individual variabilities of drug pharmacokinetics, efficacy or toxicity. For instance, incorporating uremic solute-mediated OAT1/3 inhibition was shown to improve PBPK prediction of tenofovir renal and systemic disposition^41^). Our system supports the study of disease-like fluid compositions and fluid dynamics, as well as their influence on transporters’ function and subsequent local drug PK.

The present study confirmed membrane transporter competition for creatinine, cimetidine and to a lesser extent ritonavir. Using the same setup, we were able to simultaneously investigate the transcriptomics and metabolomics modifications associated with substrate exposure and transcellular transport. We also showed that xenobiotic exposure may modulate the expression of other transporters, as well as the intracellular metabolism which may in turn affect local PK (see an example in Figure 4). These results support that mapping metabolite profiles, and interactions involving drug metabolite enzymes and membrane transporters, will improve our understanding of the interplay between RSS and (local) PK. Since they are limited to the OCT2/MATEs transporter pathway, further investigations will be required to extend such knowledge to other renal transporters of clinical and pharmacological importance,^7^ as well as to other tissues.

**Figure 4.**
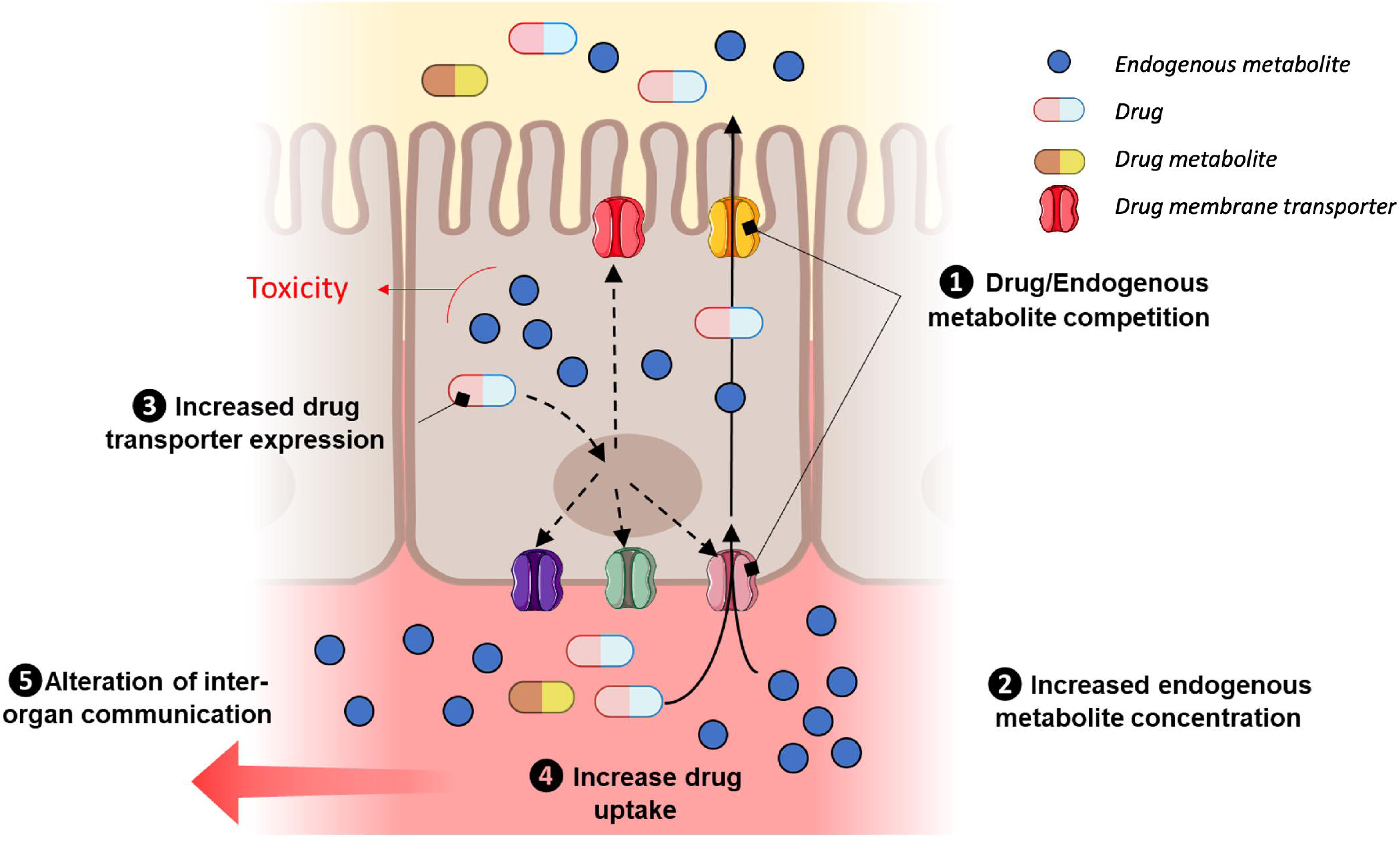
Prototypical example of transporter-mediated drug-metabolite interplay. (1) Assuming a competition (1) between xenobiotic and endogenous metabolite for influx and efflux transport, this may lead (2) to increased intracellular concentration of endogenous metabolite but also also in the systemic circulation. This might be associated with an overexpression of influx and efflux membrane transporter (3) which in turn enhance drug uptake (4) and lower systemic drug concentration. Additionally, modulation of systemic endogenous metabolite concentration will lead to transporter expression in other tissue owing to a potential alteration of RSS-mediated inter-organ communication (5).

It has been shown that ABC and SLC transporters but also drug metabolism enzymes have a distant, coordinated action over the human body, including in key organs of drug PK. For instance, network analyses of the SLC-ABC-DME relationships along the gut-liver-kidney axis confirmed the modulation of intracellular signaling pathways^38,42,43^ (*e.g.*, AHR, PXR, HNF1x and HNF4x) in distant organs. By modeling such events using multi-organ-on-chip technology, it may be possible to elucidate shared mechanism involved in either endogenous metabolite homeostasis or xenobiotic pharmacokinetics, and bridge the gap between local and systemic PK.

## 5. Conclusion

We developed a robust and reproducible kidney proximal tubule-on-chip setup able to model physiologically and pharmacologically relevant urinary excretion, which paves the way to decipher the transporter-mediated interplay between endogenous metabolite homeostasis and drug local PK. In spite of technical and experimental challenges, connecting standalone organ-on-chip devices or developing multi-organ-on-chip setups in a dynamic microfluidic environment may permit to model and better understand fundamental intracellular mechanisms and local PK events and possibly extend the remote-sensing theory to drug-endogenous metabolites interactions.

## Supporting information

Electronic Supplemental Information

## Acknowledgments

The authors thank the BISCEm platform (Univ. Limoges, Inserm US042, CNRS UAR 2015, CHU Limoges) for providing access to mass spectrometry and confocal microscopy equipment as well as Claire Carrion and Emilie Pinault for technical support.

## Supplemental information

Supplemental information are associated to the present study.

## Abbreviations

ABC: ATP-Binding Cassette
ATCC: American Type Culture Collection
DDI: Drug-Drug Interactions
DMI: Drug-Endogenous Metabolite Interactions
FSS: Flow Shear Stress
ITC: International Transporter Consortium
MATE: Multidrug and Toxin extrusion protein
OAT: Organic Anion Transporter
OCT: Organic Cation Transporter
OoC: Organ-on-chip
P-gp: P-glycoprotein
PD: Pharmacodynamics
PK: Pharmacokinetics
PT: Proximal Tubule
RPTEC: Renal Proximal Tubule Epithelial Cell
RSS: Remote Sensing Signaling
SLC: Solute Carrier

## Author Contributions

I. Petit: Investigation, Methodology, Data curation, Formal Analysis, Writing – original draft, Writing – review and editing

Q. Faucher: Investigation, Methodology, Writing – review and editing

J.S. Bernard: Investigation, Methodology, Writing – review and editing

P. Guinchi: Investigation, Methodology, Writing – review and editing

F.L. Sauvage: Formal Analysis, Methodology, Writing – review and editing

A. Humeau: Formal Analysis, Writing – review and editing

P. Marquet: Conceptualization, Funding acquisition, Resources, Validation, Writing – review and editing

N. Védrenne: Conceptualization, Investigation, Methodology, Data curation, Formal Analysis, Supervision, Validation, Writing – original draft, Writing – review and editing

F. Di Meo: Conceptualization, Funding acquisition, Investigation, Supervision, Validation, Project administration, Writing – review and editing

