## Supplementary material for "Proximal Tubule-on-Chip for Predicting Cation Transport: Dynamic Insights into Drug Transporter Expression and Function": Electronic Supplemental Information

**Supplementary Tables**

**Table S1.** List of primers

| Gene symbol | References |
|---|---|
| *ABCB1* | Hs00184500_m1 |
| *ABCC2* | Hs00166123_m1 |
| *ABCC4* | Hs00988717_m1 |
| *ABCG2* | Hs01053790_m1 |
| *SLC22A2* | Hs01010726_m1 |
| *SLC22A4* | Hs00268200_m1 |
| *SLC22A5* | Hs00929869_m1 |
| *SLC22A6* | Hs00537914_m1 |
| *SLC22A7* | Hs00198527_m1 |
| *SLC22A8* | Hs00188599_m1 |
| *SLC47A1* | Hs00217320_m1 |
| SLC47A2 | Hs00945652_m1 |
| GAPDH | Hs99999905_m1 |

**Table S2.** List of antibodies

| Target | Reference | Species | Dilution | Fluorescence |
|---|---|---|---|---|
| **P-glycoprotein** | ab235954 | Rabbit polyclonal | 1/200. | Na |
| **Na+/K+ ATPase** | ab210143 | Rabbit monoclonal | 1/500. | Alexa Fluor® 405 |
| **MATE-1** | ab92295 | Goat polyclonal | 1/200 | Na |
| **OCT2/SLC22A2** | ab242317 | Mouse monoclonal | 1/500 | Na |
| **OAT3/SLC22A8** | ab247055 | Rabbit polyclonal | 1/400 | Na |
| **Secondary antibody** | | | | |
| **Anti-Rabbit** | A11008 | Goat polyclonal | 1/1000 | Alexa Fluor® 488 |
| **Anti-Goat** | A-11039 | Chicken polyclonal | 1/1000 | Alexa Fluor® 488 |
| **Anti-Mouse** | ab175700 | Donkey polyclonal | 1/1000 | Alexa Fluor® 568 |

**Table S3.** List of the targeted 103 metabolites.

| Metabolites | | |
|---|---|---|
| 2-Ketoisovaleric acid | 2-aminobutyric acid | Cytidine monophosphate |
| Tyrosine | Serine | Cytosine |
| Histidine | 4-Aminobutyric acid | Deoxyadenosine |
| Thymidine | Aspartic acid | Deoxyadenosine monophosphate |
| 5-Glutamylcysteine | Biotin | Deoxycytidine |
| Glycyl-glutamine | Arginine | Deoxyguanosine |
| Threonine | 5-Oxoproline | D-Mannitol |
| Tryptophan | Pyruvic acid | D-Ribose |
| Folic acid | Aconitic acid | Formylkynurenine |
| Asparagine | Phenylalanine | Fumaric acid |
| Adenosine | Isoleucine | Glutathione |
| Proline | Guanosine | Glycine |
| Threonic acid | Urocanic acid | Guanine |
| Gluconic acid | Kynurenic acid | Guanosine monophosphate |
| 1-Methylhistidine | Adenine | Indole-3-acetic acid |
| Succinic acid | Leucine | Inosine monophosphate |
| Citric acid | 3-Methyl-2-oxovaleric acid | Lysine |
| Malic acid | Cystine | Methionine |
| Ornithine | Methionine sulfoxide | N-Acetylaspartic acid |
| Uric acid | Riboflavin | NAD |
| Hypoxanthine | Xanthine | Niacinamide |
| 4-Pyridoxic acid | Inosine | Nicotinic acid |
| 2-Aminoadipic acid | Pyridoxal | O-Phosphoethanolamine |
| Xanthosine | 3-aminopropanoic acid | Oxidized glutathione |
| Alanine | 3-hydroxyanthranilic acid | Pipecolic acid |
| Hexose (Glucose) | 4-Hydroxyproline | Putrescine |
| Citrulline | 5'-Methylthioadenosine | Pyridoxal phosphate |
| Lactic acid | Adenosine monophosphate | Pyridoxine |
| Uridine monophosphate | Argininosuccinic acid | S-adenosylhomocysteine |
| Pantothenic acid | Ascorbic acid 2-phosphate | Taurine |
| Kynurenine | Creatinine | Thymine |
| Choline | Cystathionine | Uridine |
| Glutamine | Cysteine | Valine |
| alpha-keto-glutarate | Cytidine | Xanthosine monophosphate |
| Glutamic acid | | |

**Table S4.** Metabolite expression measured by mass spectrometry under different flow rates.

| Mean | Static | 5 µL/min | 10 µL/min | 20µL /min |
|---|---|---|---|---|
| 2-Aminoadipic acid | 0.07 | 0.00 | 0.00 | 0.00 |
| 2-aminobutyric acid | 0.83 | 0.93 | 0.94 | 0.95 |
| 4-Aminobutyric acid | 5.37 | 6.55 | 6.43 | 6.39 |
| 4-Pyridoxic acid | 0.05 | 0.05 | 0.04 | 0.05 |
| 5-Oxoproline | 0.08 | 0.07 | 0.07 | 0.06 |
| Adenine | 0.21 | 0.20 | 0.19 | 0.19 |
| Alanine | 1.16 | 0.17 | 0.18 | 0.18 |
| alpha-keto-glutarate | 0.09 | 0.14 | 0.14 | 0.14 |
| Aspartic acid | 0.00 | 0.03 | 0.03 | 0.03 |
| Choline | 10.54 | 13.30 | 13.03 | 13.13 |
| Citrulline | 0.86 | 1.00 | 1.00 | 1.00 |
| Folic acid | 0.87 | 0.68 | 0.68 | 0.67 |
| Glutamic acid | 1.64 | 1.24 | 1.21 | 1.27 |
| Glutamine | 0.23 | 0.38 | 0.37 | 0.36 |
| Guanosine | 0.03 | 0.03 | 0.03 | 0.04 |
| Hexose (Glucose) | 0.11 | 0.15 | 0.15 | 0.16 |
| Histidine | 1.56 | 1.54 | 1.53 | 1.52 |
| Hypoxanthine | 0.00 | 0.23 | 0.23 | 0.24 |
| Isoleucine | 19.39 | 19.50 | 19.03 | 19.06 |
| Kynurenic acid | 0.30 | 0.31 | 0.30 | 0.30 |
| Lactic acid | 22.73 | 1.06 | 1.07 | 1.00 |
| Leucine | 3.69 | 4.03 | 3.97 | 3.97 |
| Malic acid | 0.06 | 0.00 | 0.00 | 0.00 |
| Methionine sulfoxide | 0.10 | 0.11 | 0.11 | 0.11 |
| Phenylalanine | 40.64 | 40.27 | 40.19 | 40.30 |
| Proline | 9.39 | 8.48 | 8.37 | 8.43 |
| Pyruvic acid | 0.52 | 0.49 | 0.50 | 0.49 |
| Riboflavin | 0.49 | 0.55 | 0.55 | 0.52 |
| Thymidine | 0.04 | 0.03 | 0.03 | 0.03 |
| Tyrosine | 11.90 | 11.07 | 11.02 | 10.99 |
| Uridine monophosphate | 0.01 | 0.04 | 0.04 | 0.04 |
| Xanthine | 0.03 | 0.00 | 0.00 | 0.00 |
| Xanthosine | 0.02 | 0.00 | 0.00 | 0.00 |

**Table S5.** Metabolite dosage on RPTEC/TERT1 under metformin or creatinine treatment compared to untreated condition. The cells are cultured in mono-channel device under 20 µL/min flow.

| Metabolite | Metformin | Creatinine |
|---|---|---|
| 2-aminobutyric acid | 0.80 | 0.97 |
| 5-Oxoproline | 0.43 | 1.61 |
| Adenine | 0.98 | 1.06 |
| Adenosine | 0.89 | 0.60 |
| Alanine | 0.84 | 0.95 |
| alpha-keto-glutarate | 0.53 | 0.75 |
| Arginine | 1.06 | 0.79 |
| Asparagine | 0.78 | 0.65 |
| Aspartic acid | 0.65 | 0.65 |
| Choline | 0.81 | 0.96 |
| Citrulline | 0.97 | 0.79 |
| Creatinine | 0.31 | na |
| Cystine | 0.95 | 0.52 |
| Folic acid | 1.09 | 1.18 |
| Glutamic acid | 0.88 | 0.96 |
| Glutamine | 0.50 | 0.76 |
| Guanosine | 0.63 | 0.70 |
| Hexose (Glucose) | 1.03 | 0.79 |
| Histidine | 0.90 | 0.68 |
| Hypoxanthine | 0.96 | 1.08 |
| Isoleucine | 0.77 | 1.04 |
| Lactic acid | 0.83 | 1.95 |
| L-Carnitine | 0.65 | 0.68 |
| Leucine | 0.67 | 1.04 |
| Methionine sulfoxide | 1.14 | 1.27 |
| Niacinamide | 0.79 | 0.95 |
| Ornithine | 0.71 | 0.38 |
| Pantothenic acid | 0.45 | 1.10 |
| Phenylalanine | 0.93 | 1.08 |
| Pipecolic acid | 0.34 | 1.32 |
| Proline | 0.82 | 0.83 |
| Pyruvic acid | 0.94 | 1.11 |
| Riboflavin | 0.97 | 1.03 |
| Serine | 0.85 | 0.64 |
| Threonine | 1.03 | 0.83 |
| Thymine | 1.15 | 0.88 |
| Tryptophan | 1.04 | 1.11 |
| Tyrosine | 1.05 | 1.09 |

### Supplementary Figures

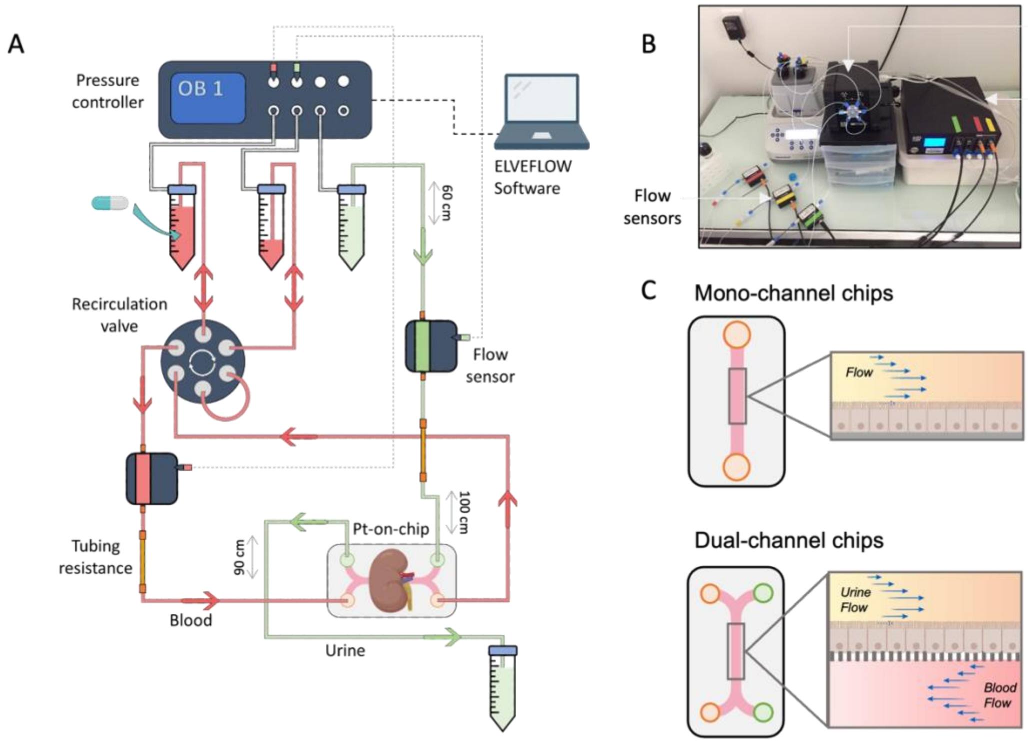


**Figure S1. Overview of commercial microfluidics setup and devise.** (A) Our microfluidic system consists of a pressure controller who will administered pressure to the system which will ensure the flow. The flow rate it's measure by flow sensors and controlled by the elveflow ™ software. The tubbing resistance permit much more precise control of the flow rates.(B) lab view of the microfluidic system. (C) Schematic view of commercial mono (ibidi ™) and dual (beonchip ™) channel device use for RPTEC/TERT1 culture.

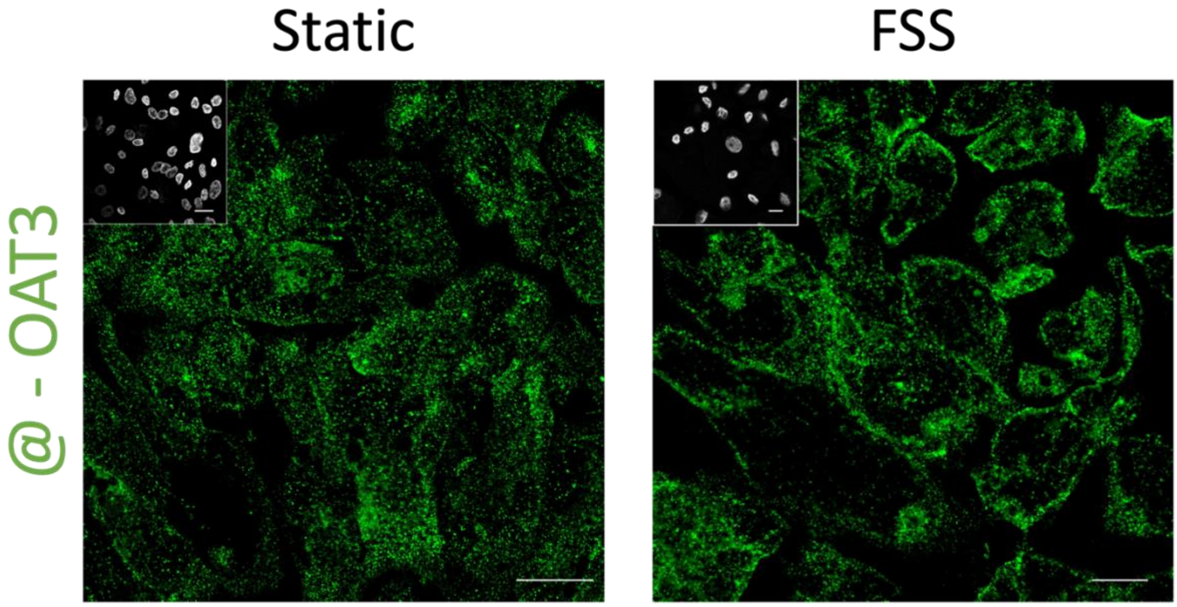


**Figure S2. Immunolabeling of OAT3 on RPTEC/TERT1.** Cells are cultured under undetermined flow condition or FSS (20 µL/min flow; 0.02 dyn/cm²) condition. Insert represent DAPI staining, scale bar represents 20µM.

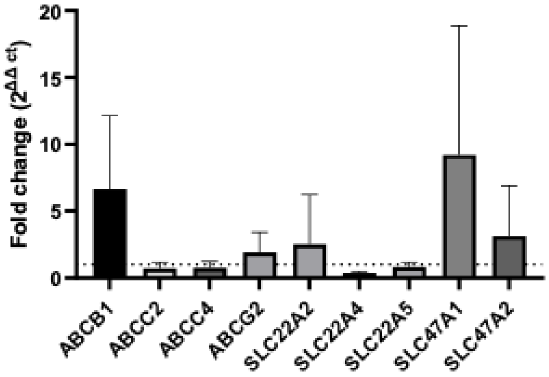


**Figure S3. Fold changes regarding the relative expression levels of mRNA transporter expressions of RPTEC/TERT1 cell line in dual-channel** device under 10 µL/min (0.03 dyn/cm²) in apical compartment and 20 µL/min (0.07 dyn/cm²) in basal compartment (N=4). Error bars represent standard deviations. Statistical significance was determined using Kruskal Wallis test, asterisks (*) indicate statistically significant differences with respect to mono-channel device under 20 µL/min (0.02 dyn/cm², N=4), *p-value < 0.05, **p-value < 0.01, ***p-value < 0.001).

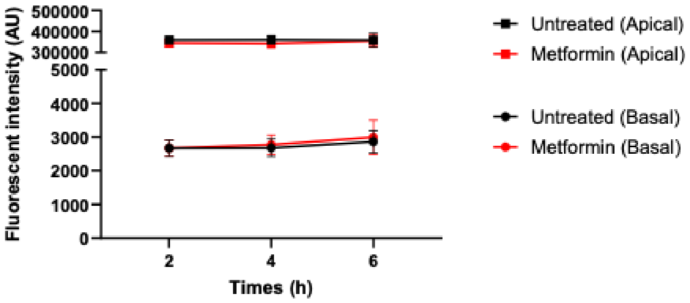


**Figure S4. FITC-dextran permeability assay.** The permeability of the cell layer was assessed by measuring the rate of fluorescein isothiocyanate (FITC)-dextran passage in presence or not of metformin 100µM, from the apical to the basal compartment of a transwell system. 20µM of FITC-conjugated 10kDA dextran (FITC-dex, Sigma-Aldrich) was introduced into the apical compartment and the FITC-dextran transport across the barrier was determined by serially sampling fluid from the apical and basal compartment and measuring its fluorescence intensity, which was used as an index of barrier permeability.

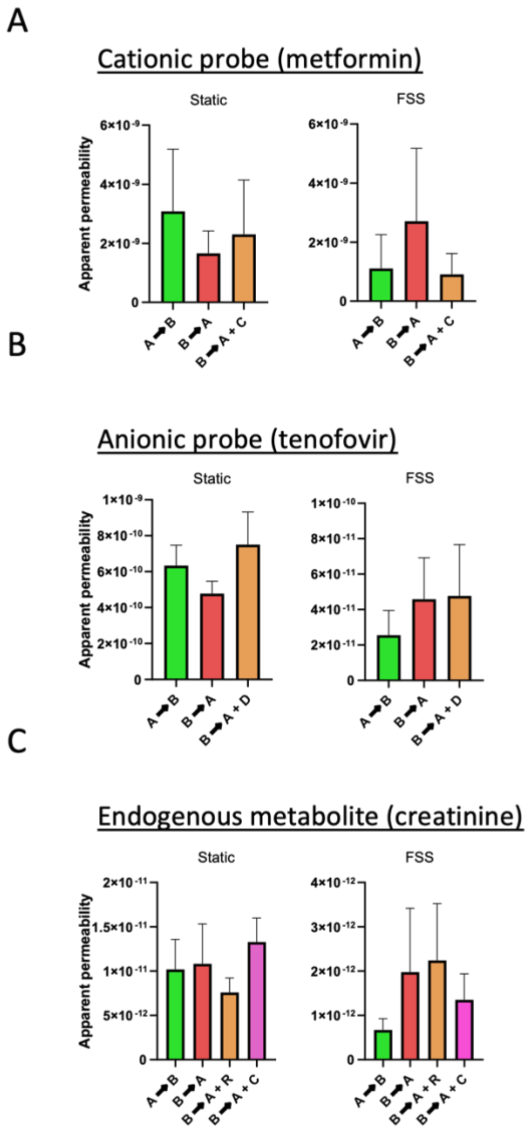


**Figure S5. Calculated apparent permeabilities for transcellular transport.** Apparent permeabilities were calculated for (A) metformin (100 μM, N=4), (B) tenofovir (30 μM, N=4) and (C) creatinine (10 μM, N=8) as cationic, anionic and endogenous probes, respectively. $P_{app}$ were calculated considering transcellular transport from apical to basal compartment (A→B) and basal to apical compartment (B→A) for static and FSS conditions in presence or not of an inhibitor, namely cimetidine (C, 100 μM, N=3) for metformin; diclofenac (D, 30 μM, N=4) for tenofovir and ritonavir (R, 15 μM, N=3) or cimetidine (C, 100 μM, N=3) for creatinine.
